# Integration of transcriptomics data into agent-based models of solid tumor metastasis

**DOI:** 10.1101/2023.01.09.523238

**Authors:** Jimmy Retzlaff, Xin Lai, Carola Berking, Julio Vera

**Affiliations:** Department of Dermatology, Universitätsklinikum Erlangen and Friedrich-Alexander-Universität Erlangen-Nürnberg, Laboratory of Systems Tumor Immunology, Erlangen, DE; Deutsches Zentrum Immuntherapie, Erlangen, DE; Comprehensive Cancer Center Erlangen-EMN, Erlangen, Bayern, DE

## Abstract

Most of the recent progress in our understanding of cancer relies in the systematic profiling of patient samples with high throughput techniques like transcriptomics. This approach has helped in finding gene signatures and networks underlying cancer aggressiveness and therapy resistance. However, -omics data alone is not sufficient to generate insights into the spatiotemporal aspects of tumor progression. Here, multi-level computational models are promising approaches, which would benefit from the possibility to integrate in their characterization the data and knowledge generated by the high throughput profiling of patient samples.

We present a computational workflow to integrate transcriptomics data from tumor patients into hybrid, multi-scale models of cancer. In the method, we employ transcriptomics analysis to select key differentially regulated pathways in therapy responders and non-responders and link them to agent-based model parameters. We next utilize global and local sensitivity together with systematic model simulations to assess the relevance of variations in the selected parameters in triggering cancer progression and therapy resistance. We illustrate the methodology with a *de novo* generated agent-based model accounting for the interplay between tumor and immune cells in melanoma micrometastasis. Application of the workflow identifies three different scenarios of therapy resistance.

## 1 Introduction

In the last decade we have progressed remarkably in our understanding of cancer pathogenesis and metastasis, and this has had positive consequences in our ability to diagnose, stratify and treat metastatic tumors. In line with this, immune evasion is a hallmark of metastatic cancer (Hanahan and Weinberg 2011) and our deep understanding on its mechanisms of action has been vital to the development of therapies such as immune checkpoint inhibitors (ICI) for aggressive tumors like advanced melanoma (Wei et al. 2018), which greatly increased patient survival in metastatic melanoma patients (Larkin et al. 2015). A significant fraction of the progress in cancer research is due to the characterization of tissue samples from large cohorts of patients through genomics, transcriptomics, proteomics and/or epigenomics analysis. These techniques give access to quantitative data describing the activation and expression of (all) genes in cancer, thereby providing the information necessary to investigate the genetic landscape of cancer progression (Akbani et al. 2015) and to reconstruct and dissect the gene regulatory networks underlying cancer pathogenesis and therapy response (Dreyer et al. 2018). However, -omics data alone cannot account for some relevant levels of (de)regulation happening in cancer, which are linked to spatiotemporal variations in the tumor’s molecular and cellular composition, as well as to the existence of nonlinear regulatory structures like feedback and feedforward loops (Lai et al. 2016).

In this context, mathematical modeling and in particular multi-level spatial computational models are a viable method as they allow to investigate the dynamic behavior of the tumor microenvironment (TME) from hypothesized cell behavior and to evaluate therapeutic strategies (Metzcar et al. 2019). These models can describe and simulate the dynamics of cancer-deregulated intracellular gene circuits (Kirouac et al. 2017), but they can also be used to integrate genes and gene circuits activity into tissue-scale models of cell-to-cell interactions (Vera et al. 2013).

In such agent-based models (ABM), cells act as discrete individuals according to their set of rules. ABMs of cancer immune environments have been recently reviewed by Norton et al. (2019). There are some challenges to devising these computational models in the context of cancer: First, there is a trade-off between detailed modeling of biological features with different scales occurring in cancer progression and keeping the model simple enough to allow interpretability and reasonable computing effort. Second, these model types include a large amount of model parameters that require diverse, quantitative data to be characterized. In the case of tissue modeling, many parameters can only be calibrated indirectly, as the experimental modalities to observe single cell behavior *in vivo* are missing. We also need domain knowledge to decide which parameters are relevant for model simulations and investigating hypotheses, allowing to propose or optimize therapies.

In this paper, we propose to integrate transcriptomics analysis from cancer patient cohorts in the design and characterization of agent-based model simulations. To this end, we describe a method for analyzing transcriptomics data, ranking and selecting key gene sets underlying a condition of interest and linking these to selected agent-based model parameters. We utilize this information to prioritize parameters for investigation of the model behavior, as an analysis of the whole parameter space is computationally infeasible., The influence of these priorized parameters is investigated via global sensitivity analysis and massive, systematic model simulations. We exemplify the use of the method for a case study on melanoma metastasis and immunotherapy resistance. To this end, we built an agent-based model accounting for the interplay between tumor and immune cells in a micrometastasis.

## 2 Materials and Methods

In this work we followed a workflow sketched in Fig. 1. It contains several steps including model construction, exploration and calibration, linking enriched gene sets to parameters, and analysis. The steps we take are discussed in detail below.

**Fig. 1.**
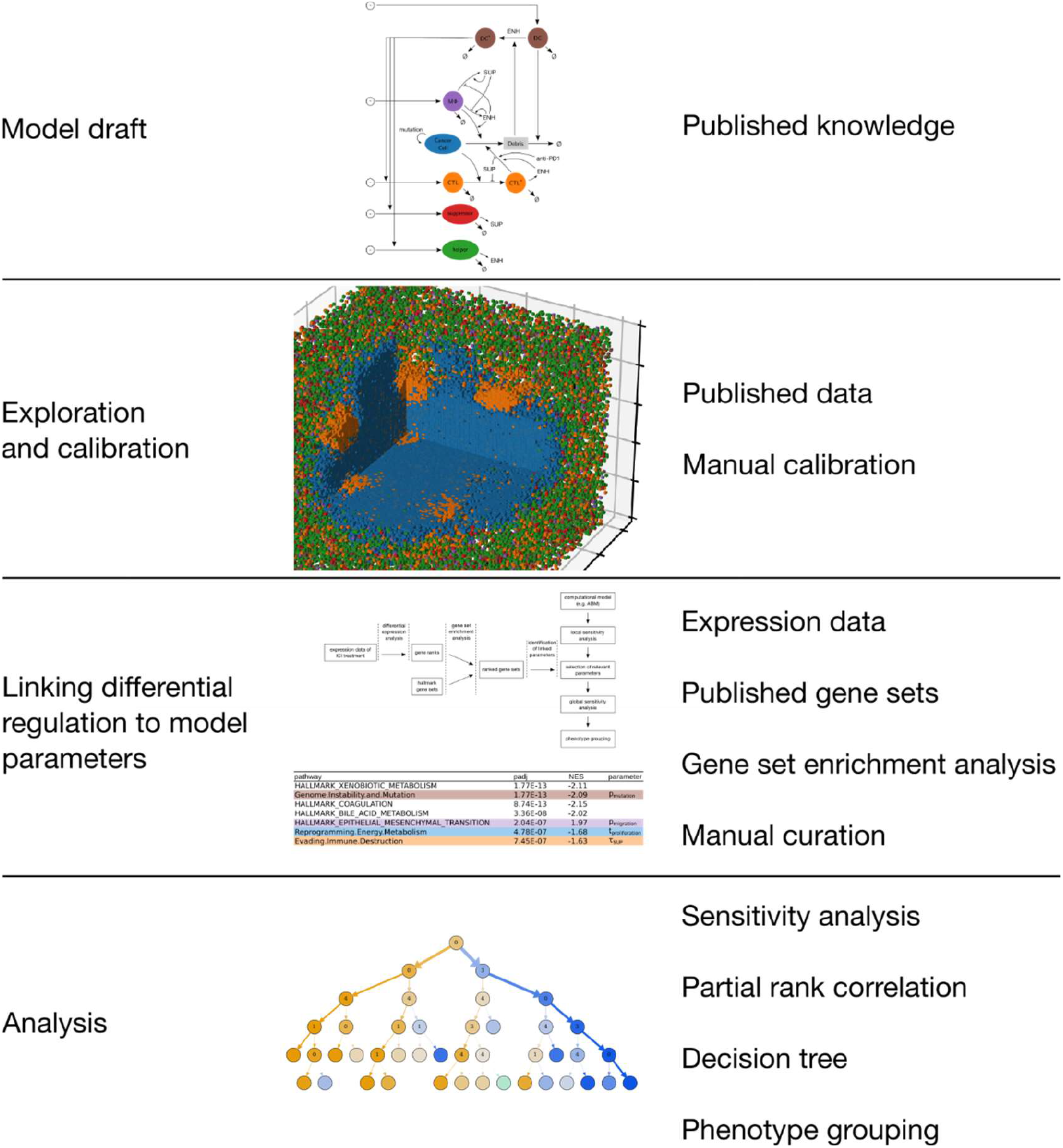
Workflow of the study. On the left are labels of the four overarching steps, on the right is a brief list of the central materials and methods used in each step. We first created the model and set a nominal parameter configuration. Then we linked expression data to model parameters to narrow a selection of parameters whose influence we analyzed in more depth.

### 2.1 Multi-level melanoma immunology model

The general modeling concept we used is to create a spatial agent-based model of the TME that interacts with a systemic compartment. This is a modeling approach that has been suggested in recent reviews (Norton et al. 2019; Metzcar et al. 2019) and was followed in other multiscale models as well (Santos and Vera 2020; Gong et al. 2017). We built our model based on knowledge of tumor immunology and signaling in cancer and melanoma (Abbas et al. 2014; Marzagalli et al. 2019). The model contains immune and tumor cells in the melanoma TME and their interactions, which can be either based on cell-cell contact or on intercellular communication through cytokines.

The model accounts for parts of the innate and adaptive immunity to tumors including immunosurveillance without considering memory and long-term immunity. For the immunosurveillance we assumed that the tumor antigens are not yet detected by the adaptive immune system. A sketch of the model and in particular the considered cell interactions are shown in Fig. 2. Specifically, the immune cells accounted for are cytotoxic T lymphocytes (CTLs), T helper cells (Th), B cells, regulatory T cells (Tregs), dendritic cells (DCs), macrophages and myeloid-derived suppressor cells (MDSCs).

**Fig. 2:**
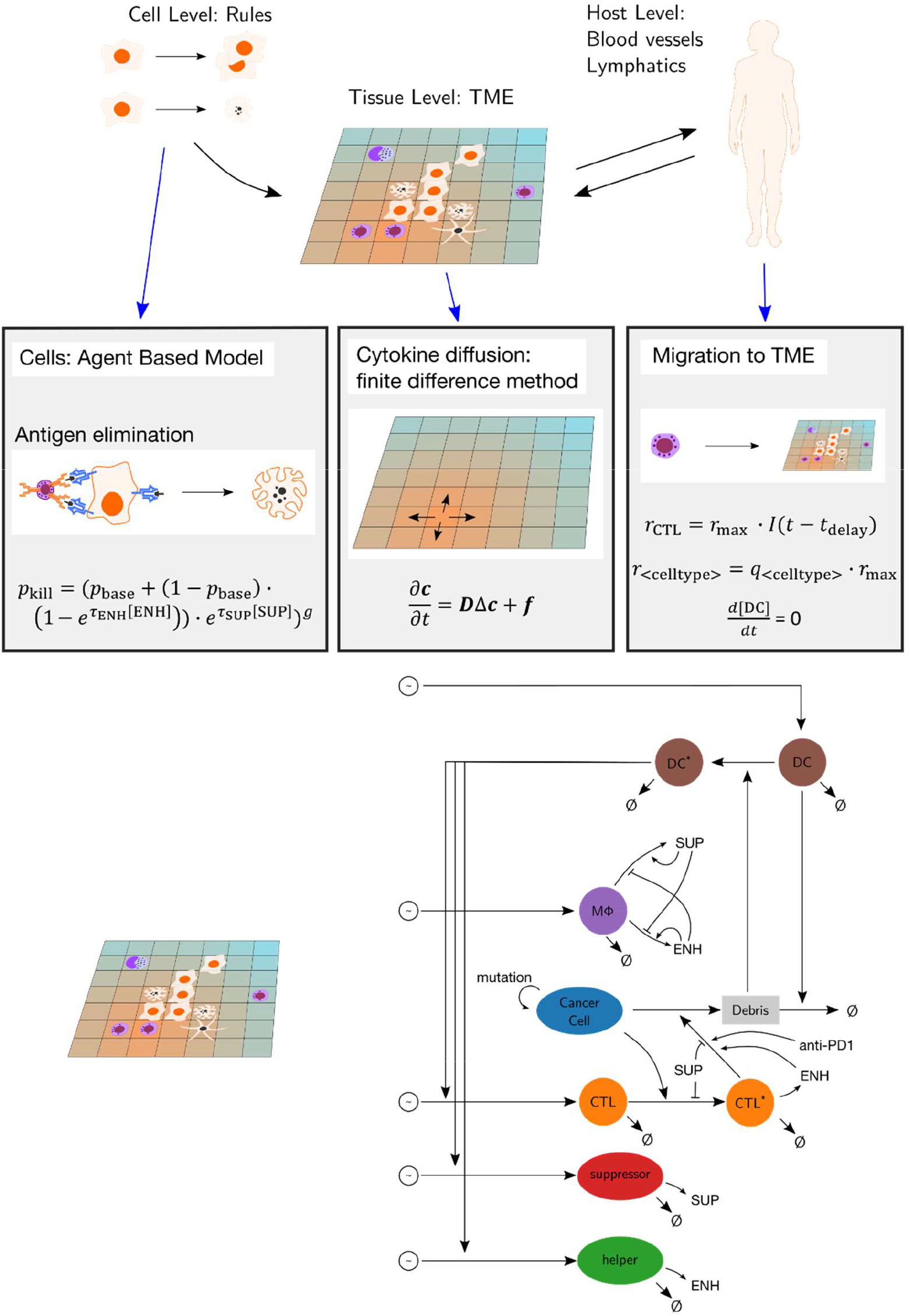
Concept of the model. On the cellular level, an agent-based model is used, where cell physiology is described as cell type specific rules. This is coupled on the level of the tumor-microenvironment with the cytokine diffusion solver and the recruitment model of immune cells. Cancer cell debris is detected by DCs which will attract CTLs, suppressors and helpers after a delay. CTLs that contact cancer cells switch to an activated state and can kill the cancer cell. The killing probability is influenced by cytokines and applied anti-PD1 therapy. Abbreviations: ENH: immunoenhancing cytokine, SUP: immunosuppressive cytokine, DC: dendritic cell, MΦ: macrophage, CTL: cytotoxic T lymphocyte.

We assumed that the communication between immune cells is mediated by cytokines and chemokines. The cell behavior is modeled as logical rules. Further, our model includes helper cells and suppressor cells as abstract cell types that account for immune cells that primarily have a regulatory role, such as CD4+ T cells, B cells and MDSCs. These cells influence the immune response via secreting cytokines.

We labeled the involved cytokines based on whether they have a primarily immunoenhancing or immunosuppressive effect and modeled two abstract surrogate cytokines accordingly: immunoenhancing (ENH) and immunosuppressive cytokine (SUP). ENH accounts for cytokines that increase the effectiveness of cytotoxic mechanisms like IFN-γ, as well as for chemoattractants for cytotoxic cells such as CCL3, CCL4, CCL5, CXCL9 or CXCL10. Examples for molecular species that have immunosuppressive effects are IL-10, TGF-β and IDO. The surrogate cytokines keep the model simpler by assuming that the cytokines do not have pleiotropic effects, although it has been shown that some cytokines may trigger both immunosuppressive and immunoenhancing effects (Donia et al. 2016). For instance, IFN-γ, a key regulator of the adaptive immune response, can trigger both the expression of major histocompatibility complex I (MHC-I) and of PD-L1. The former increases the recognition of cancer cells by T cells, while the latter inhibits the effector mechanism of T cells.

#### 2.1.1 Cytokine diffusion

Cytokines are modeled using a continuum model that tracks cytokine concentrations rather than discrete molecules. Their diffusion is described by Fick’s second law

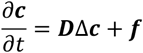

with the compounds’ concentrations ***c***, the diffusion constants ***D***, and a term ***f*** accounting for secretion and degradation. Cytokine diffusion is solved using a finite difference method with the Euler forward method.

#### 2.1.2 *In situ* cell populations

##### Cancer cells

A cancer cell population of 125 cells is seeded at the lattice center at the beginning of a simulation. As the tumor cells migrate and divide, they will spread over the tissue in the course of a simulation.

Cell motility is implemented as a random walk, allowing a cell to move to neighboring lattice positions with a probability *p*_migration_. If the chosen position is already occupied by another cell, the cells either swap places or stay at their positions with equal probability. In general, it is assumed that cancer cells are less motile than immune cells.

Cancer cells can die with a probability *p*_death_, which leads to their removal from the lattice. Dead cancer cells will leave debris that can be collected by dendritic cells and facilitate an immune response.

Cancer cells are considered to have uncontrolled replication potential and will attempt to divide after a fixed length of time *t*_proliferation_ has passed, which accounts for cell growth and cycle. Dividing cells are temporarily immobile for the time step where the cell division occurs. Cell division can only take place if there is a vacant neighboring position that a daughter cell can occupy. This rule implicitly models cell contact inhibition, a trait that cancerous cells usually lose (Hanahan and Weinberg 2011). However, this assumption is in line with previous studies that use lattice-based models (e.g. Wang et al. 2013). It simplifies the modeling of cell mechanics, and evades calculations such as equilibria of forces between cells and the consequential possibility of cells pushing each other away.

Cancer cells may present one or multiple tumor-specific antigens depending on their mutations. We modeled only passenger mutations, meaning if a cancer cell mutates, it may start to present another antigen. This can induce an adaptive immune response specific to that antigen. Cancer cells mutate at each time step with a probability *p*_mutation_. The mutations that are modeled are non-driver mutations, as they only affect the antigen pool that a particular cancer cell presents, which influences its susceptibility towards clearance by CTLs: more mutations lead to a larger antigen pool and recognition by different CTL clones. The mutations are modeled as a finite allele model, with 32 possible mutations. This allows to track the individual mutations, but has the disadvantage that it is not realistic compared to the near-infinite mutations possible in a real human genome, potentially leading to artifacts, e.g. through the possibility of reverse mutations.

##### Cytotoxic T lymphocytes

Cytotoxic T lymphocytes patrol the tumor site and are able to induce apoptosis in cancer cells upon contact. A cell is considered to be in contact with another if it is present in its Moore neighborhood (i.e. adjacent cell including diagonal adjacency). A CTL recognizes an antigen that is specific to its receptor and kills cancer cells presenting the antigen. It probes its neighborhood for a recognizable cancer cell in random order. The randomness is introduced to avoid a direction bias that might lead to simulation artifacts. If the CTL recognizes a cancer cell, it kills it with a probability modeled as

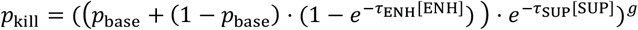

that depends on a base killing probability *p*_base_, the local concentrations of SUP [SUP] and ENH [ENH], respective rate constants *τ*_SUP_ and *τ*_ENH_, and the influence of anti PD1 checkpoint inhibitor therapy *g*. The probability is modeled in such a way that killing of tumor cells by CTLs becomes more effective in the presence of ENH and less effective in the presence of SUP. Anti PD1 therapy is modeled as a power law influence (Vera et al. 2007) and is set to 1 (no therapy) and can be toggled to 0.1 during a simulation (application of therapy). At this abstraction level of the model it is indifferent whether the drug targets PD1 which may be expressed by CTLs or its ligand PDL1 which may be expressed by the cancer cells, as it only influences the killing mechanism on contact. The killing has a duration *t*_kill_, in which the CTL becomes immobile and unable to kill other neighboring cancer cells.

CTLs undergo apoptosis after a fixed life span *t*_life,CTL_ expires. Note that CTL expiration accounts for different cell fates including exhaustion, apoptosis or leaving the TME. We do not model CTL proliferation in the TME, although it has been reported in cases of combination therapy (Spranger et al. 2014). Unlike cancer cells, a CTL follows its migration rule at every time step unless it is in an immobile state. Therefore, its motility depends only on the cell density in its vicinity. It is capable of performing both random walk and chemotactic migration, following the ENH gradient. By default, CTLs perform random walk and they change to chemotactic migration if the concentration of surrounding ENH cytokines exceeds a threshold. We modeled this threshold to prevent the CTLs from being sensitive to very low ENH concentrations. The CTLs moving in the chemotactic mode are in an activated state. The activated CTLs secrete ENH cytokines with a fixed rate *r*_ENH_. ENH increase their cytotoxic capabilities and attract other CTLs to their vicinity, allowing fast finding and clearance of cancer cell colonies.

##### Dendritic cells

DCs are antigen-presenting cells to immune effector cells such as T cells. In the model, DCs function as probes for tumor cells, and they move inside the lattice at each simulation step if a vacant position in the vicinity is available. We did not consider apoptosis of DCs in the TME, as we assume that DCs are either tissue-resident or filtrated through the TME during their lifetime. A DC will collect all cancer cell debris it encounters. It then starts to present the antigens it processed. Furthermore, it becomes activated and leaves the tumor site. Once a DC has left the tumor site, it increases a signal that leads to a delayed recruitment of CTLs that are specific to the antigens it now presents. This way we implicitly modeled the homing of DCs to the tumor-draining lymph node. We assume homeostasis of the DC population, and for every DC that leaves the tumor microenvironment, a new DC will be recruited.

##### Helper and Suppressor cells

Helper cells account for CD4+ helper T cells as well as tumor infiltrating B cells. They constantly secrete ENH cytokines (*r*_ENH_). Suppressor cells primarily account for regulatory T cells (Treg) and myeloid derived suppressor cells (MDSCs). Analog to helper cells, they constantly secrete SUP cytokines. Both helper and suppressor cells perform a random walk during their lifetime, which is fixed to *t*_life,helper_ and *t*_life,suppressor_, respectively.

##### Macrophages

Macrophages have cytotoxic capabilities, and can secrete both ENH and SUP cytokines. Similar to CTLs, their cytotoxicity is influenced by cytokines. Their cytokine secretion rates depend on the ratio of the concentrations of local SUP and ENH cytokines:

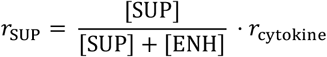

and

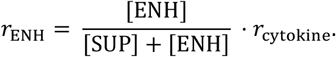

The equations result in positive feedback loops, making the secretion rates of SUP and ENH by macrophages positively correlate with their own concentrations. The feedback loops imitate an environment-dependent phenotype plasticity that resembles M1 and M2 phenotype activation described in the literature (Kim and Bae 2016).

#### 2.1.3 Cell recruitment

Newly recruited cells appear on a free position at the border of the TME lattice based on the assumption that recruited cells enter from nearby blood vessels. The cell types recruited to the TME are CTLs, DCs, macrophages, helper and suppressor cells.

The recruitment of CTLs is preceded by DC-induced clonal expansion and differentiation in the lymphatic tissues, which introduces a delayed response. Therefore, CTLs are recruited to the TME with an antigen specific rate *r*_CTL_ that depends on delayed tumor detection of DCs.

The delay is modeled as a queue with a fixed size *t*_delay_ for each antigen. At each simulation step, the oldest value will be dequeued and leads to recruitment of CTLs, while a new value will be enqueued and initialized to zero. Every DC presenting the respective antigen that leaves the tumor site at the simulation step will add a number to the new value in the queue, leading to immune cell recruitment in the future. Helper cells are recruited alongside CTLs with a fixed ratio of 1:1, which approximates reported data (Hernberg 1996).

Recruitment of suppressor cells and macrophages depend on CTL recruitment with fixed ratios *q*_<cell type>_ e.g.

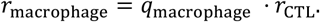

### 2.2 Model environment, simulation and parameterization

The lattice is modeled with cubic cells with a side length of 10μm, with 100 × 100 × 100 cells, representing a volume of 1mm^3^. We set *Δt* = 10 min for the duration of a simulation step, which is taken from a similar model by Gong et al. (2017). The fastest action a cell undertakes and that is affected by the step size is cell movement. The time step and cell length correspond with a maximum cell speed of 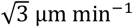. The maximum cell speed is therefore about 10 times slower than the speed of neutrophils performing chemotaxis in a microfluidic device and about 5 times faster than H69 small cell lung cancer cells (Milo et al. 2010). We assume this depicts cell motion with an adequate speed assuming that leukocytes move more slowly in tissue than in a microfluidic device. Simulations start with a small homogeneous population of 125 cancer cells at the center of the lattice and random uniformly distributed populations of DCs and macrophages. We run simulations for a period of 100 days or 8401 steps, respectively. This period is chosen on the assumption that a successful immune response will clear the metastasis within 50 days, as is indicated for adaptive immune responses (Abbas et al. 2014). We doubled the simulation time to investigate the model progression of small residual cancer cell populations that many simulations showed at day 50. Further, a simulation will abort earlier if the cancer cell population grows larger than 800,000. This abortion condition is chosen to limit the computational effort of the simulations. We consider it justified as 80% of the lattice spaces will be occupied by cancer cells, effectively simulating a tumor expansion beyond the model space.

We used published experimental data to calibrate as many parameters as possible, whose annotation and nominal values are listed in supplementary Table S1. To determine the maximum recruitment rates, we use ratios of cell types that are described in literature, leaving but one recruitment rate uncharacterized. With this modeling choice we achieve that the simulated immune infiltrate resembles an infiltrate found in experiments over the course of a simulation.

Running a single simulation took about 1 hour and 20 minutes on our hardware (4 Intel Xeon E5-4660, 256GB RAM), requiring about 10 MB RAM. For the sensitivity analyses, we ran up to 64 simulations in parallel. We implemented our model in c++17, using HDF5 and json data formats for I/O. To account for the stochasticity in the model, we repeated each simulation multiple times with different seeds for the random number generator. To decide on a number of replicas to make for each simulation, we ran simulations with the nominal parameter configuration both with and without application of ICI therapy and compared the 95% confidence intervals of the expected cancer cell populations after 3 and 100 replications, and found that using a low number of replications is acceptable for our analysis (supplementary Fig. S1). For the local sensitivity we repeated each simulation 10 times, and for the global sensitivity analysis 3 times.

### 2.3 Linking differential regulation to model parameters

For computationally expensive model simulations, global sensitivity analysis is only feasible for a small subset of model parameters. Here we propose to base the selection on parameters linked to biologically relevant gene sets using a workflow as depicted in Fig 3. We performed a gene set enrichment analysis to identify and characterize a subset of them that is related to response to anti-PD1 treatment. First, we downloaded and processed the transcriptome data (GSE78220) of pre-treatment melanomas undergoing anti-PD1 checkpoint inhibition therapy. Second, we identified differentially expressed genes between responders and non-responders. Third, we performed a gene set enrichment analysis using the differentially expressed genes and identified gene sets in which the genes are involved. We assumed that the identified enriched gene sets are crucial for the pathogenesis and progression of melanoma, and therefore we manually annotated them with corresponding model parameters.

**Fig. 3.**
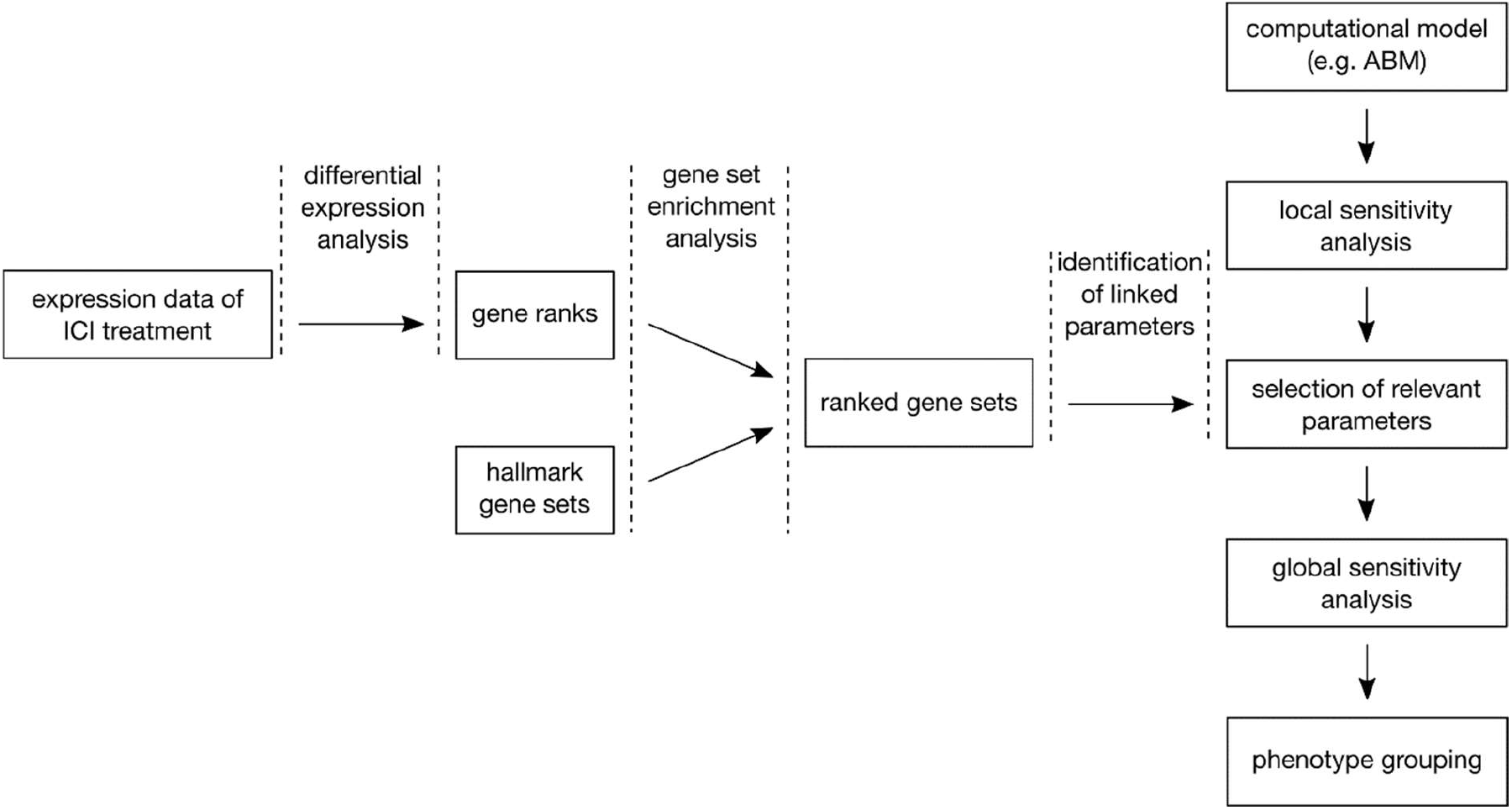
Method of linking expression data to model parameters. Besides performing a local sensitivity analysis to preselect a set of parameters for global analysis, we propose to enrich expression data of different conditions to link them to parameters of potential biological relevance.

For the gene set enrichment analysis, we used the R package fgsea (Sergushichev 2016) that tested the enrichment of the identified differentially expressed genes using the MSigDB hallmark gene set collection (Liberzon et al. 2015) and cancer hallmark genes (CHG, Zhang et al. 2020). The fgsea algorithm searches for gene sets where highly ranked genes are enriched. It is given a ranked list of genes and a list of gene sets. We calculated gene ranks based on differential expression as

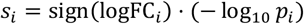

with the binary logarithmic fold change logFC_*i*_ and the p-Value *p*_*i*_. To explore the enrichment, we examine the Benjamini-Hochberg adjusted p-value and list the significantly regulated gene set. These were manually annotated with model parameters they relate to, excluding those that could not be related (e.g. because of their generality or association to processes that are not modeled).

Next, we selected parameters for global sensitivity analysis based on the gene set enrichment and the local sensitivity analysis. In our case, we could afford to run a global sensitivity analysis for 5 parameters, while 6 parameters were identified with the gene set enrichment. To exclude one parameter from this selection, we performed a local sensitivity analysis (i.e. one at a time perturbation) and exluded the least influential parameter

### 2.4 Sensitivity analysis and decision tree based phenotype grouping

We selected five model parameters of interest for sensitivity analysis, which we set up in a quasi Monte Carlo fashion where we sample the selected parameter space using the Sobol’ sampling sequence implementation of chaospy (Feinberg and Langtangen 2015). As boundaries for the parameter space we set (0, 2) times the nominal value. Assuming that about five simulations per parameter are needed to sufficiently cover the parameter space, we sample 5^5^ = 3125 parameter sets. As the model includes stochastic processes such as killing or moving probabilities, we repeat each simulation 3 times, leading to a total number of 9,375 simulations for the sensitivity analysis.

To analyze the sensitivity of our parameter selection, we trained a decision tree as a meta-model, with the aim to find sensitive parameter subspaces and the parameter sensitivities on the simulation outcome (Hastie et al. 2009; Saltelli et al. 2008), a similar approach has been followed in an earlier work (Santos et al. 2018). To quantify the parameter sensitivities, we used partial rank correlation (Marino et al. 2008) and feature importances derived from the decision tree. As target variable we chose the cancer cell population at the end of the simulation, labeling the simulation results either as emerging metastasis (>700,000), complete remission (0) or else residual disease. This gives an indication of how good the immune response is in eliminating the emerging metastasis. There are some limitations though, as we can generally not assume that a simulation will reach a steady-state by its end.

Decision trees have the advantage that they can reproduce nonlinear and non-monotonous behavior, which is useful as we cannot assume a linear model behavior a priori. We use the scikit-learn implementation, which also calculates normalized parameter importances on the regression splits (Pedregosa et al. 2011). As an optimization criterion we chose the Gini impurity. The simulations were randomly split into a training and test data set (80:20) and 5-fold cross-validation was carried out on the training set yielding a mean accuracy of 0.9 and a standard deviation of 0.004. To avoid overfitting and to keep the decision tree easily human-interpretable we constrain it to a depth of 5 and a minimum split size of 3.3%.

## 3 Results

We developed a model of the immune reaction to melanoma that aims to account for the core cellular mechanisms influenced by the cytokine milieu. The TME is set up to simulate a newly seeded micrometastasis, where a cancer cell colony grows, is detected by immunosurveillance and challenged by both innate and adaptive immune responses, where CTLs are the cytotoxic actors.

### 3.1 Selection of the nominal model configuration

We calibrated most model parameters to data estimates from the literature (supplementary Table S1). To find values for the three parameters to which no data was found, we explored their parameter space to find a sensitive parameter set that we fixed as nominal values (supplementary Fig. S2).

In Fig. 4 we show simulations of the nominal parameter configuration and a configuration with reduced recruitment of immune cells with and without ICI treatment. This is motivated by findings that immune cell infiltration or CTL infiltration in particular is correlated with ICI treatment outcome (Li et al. 2021, Kümpers et al. 2019, Du et al. 2021, Nie et al. 2019), which we tested if our model would replicate. Model parameters have been randomly perturbed within the range of +/-25% to account for patient diversity. It can be seen that for the nominal parameter configuration, simulations without ICI lead to emerging metastases in almost any case. Simulations with low CTL infiltration and application of ICI lead to remission in 76/100 and to residual disease in 24/100 simulations. Simulations with ICI and the nominal (high) CTL infiltration lead to complete removal of cancer cells in 99/100 simulated cases and residual disease in one case. Taken together, the selection of parameter values for the nominal model configuration renders results that qualitatively match clinical evidence of patient response with high and low CD8 cell infiltration (Li et al. 2021, comp. Fig. 4C).

**Fig. 4.**
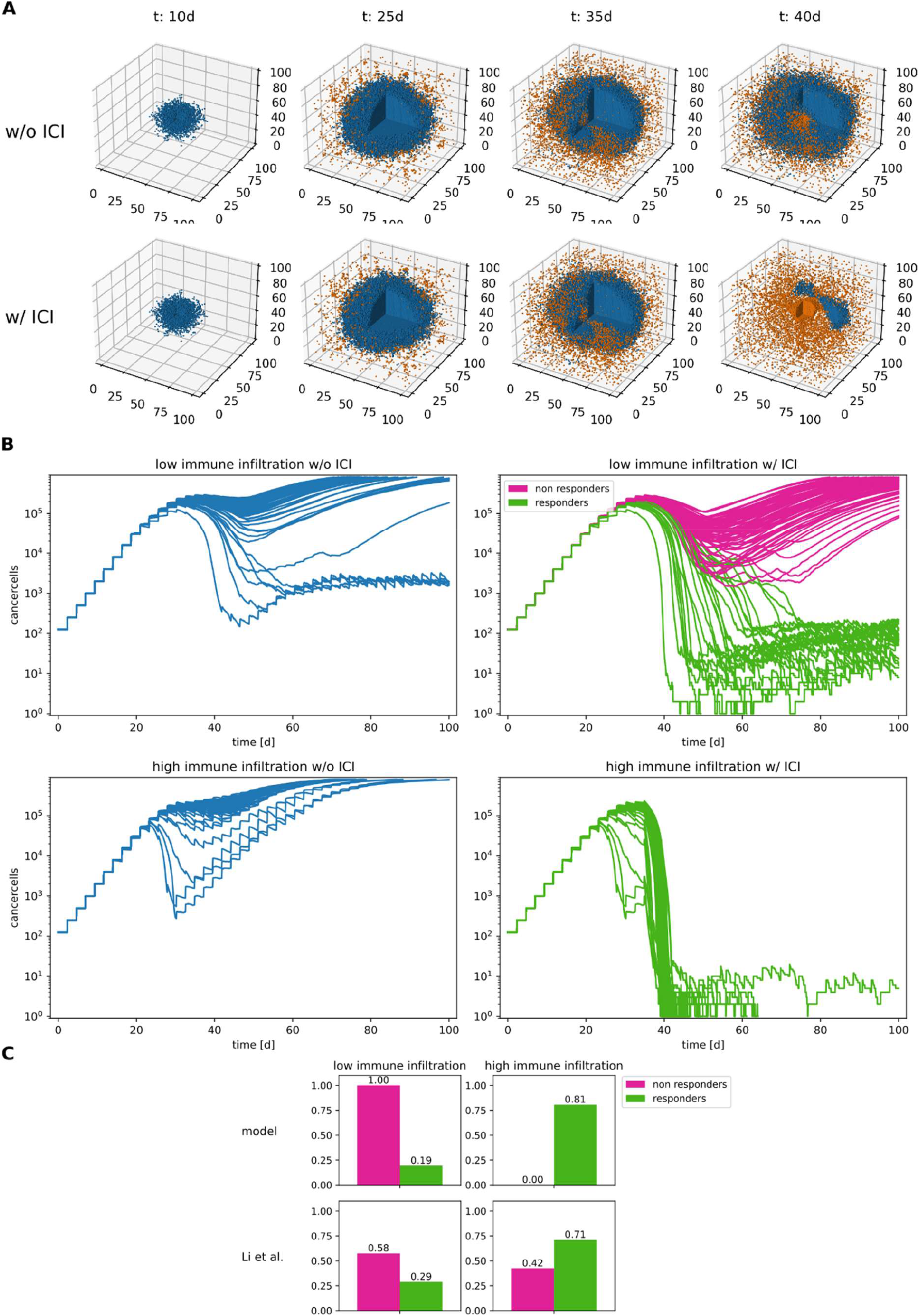
Simulations of the nominal parameter configuration with and without ICI therapy. A: comparison of the cell lattices over the course of a simulation. Only cancer cells (blue) and CTLs (orange) are shown for clarity B. Cancer cell populations of 100 simulations per condition with randomly perturbed model parameters (+/-25%). High immune infiltration marks the nominal configuration, low immune infiltration simulations have a 3-fold reduced immune cell recruitment rate. The simulations with ICI are designated “responders” or “non-responders” depending on their final cancer cell population. C. Qualitative comparison of the conditions with fractions of anti-PD1 responders and non-responders with high and low CD8 infiltrates respectively.

### 3.2 Transcriptomics data-driven selection of therapy-response related gene sets and their connection to model parameters

Gene set enrichment analysis using differential gene expression (anti-PD1 responders vs. non-responders) resulted in 24 significantly differentially regulated gene sets (Benjamini-Hochberg adjusted p-value <= 0.05). Fig. 5 demonstrates how we used this data to identify model parameters that are of particular interest, which we selected for the computationally expensive global analysis. For the parameter selection we considered the significantly differentially regulated gene sets. We annotated and mapped them to the corresponding model parameters. We excluded 14 gene sets that could not be linked to any model parameter either because they are generic, disease-specific, or not directly related to any modeled mechanism.

**Fig. 5.**
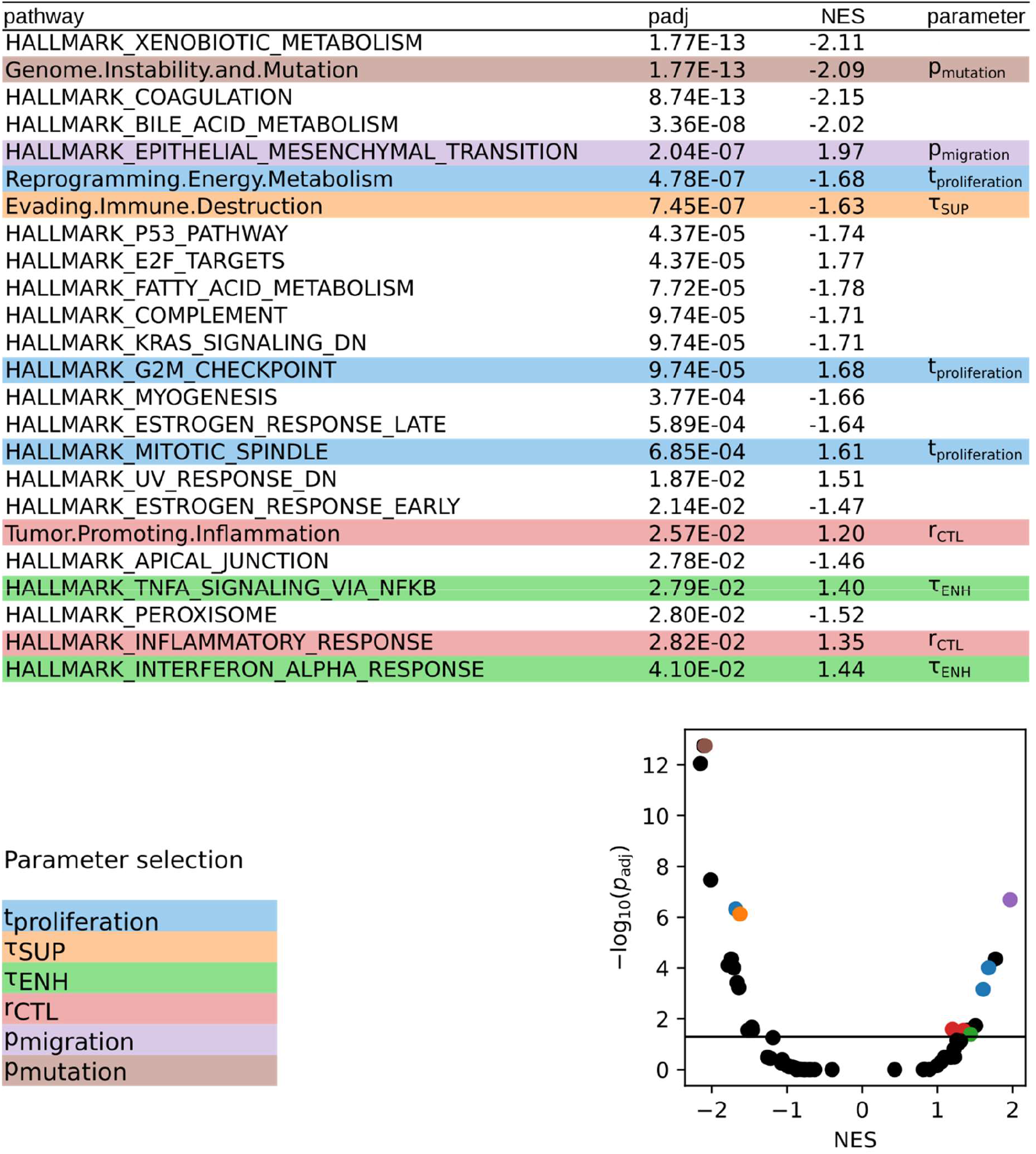
Central step of the proposed method to select model parameters that are related to differential regulation in benign vs. malignant tissue. Top: Significantly differentially regulated gene sets in melanoma sample of different ICI treatment response that could be linked to model parameters. See full list in supplementary Table 2. Bottom left: List of the identified parameters connected to the selected gene sets. Bottom right: volcano plot of the adjusted p value against the normalized enrichment score. Gene sets accounting for parameters are marked with corresponding colors.

We link one gene set, “genome instability and mutation”, to the mutation probability *p*_mutation_. Another, “epithelial-mesenchymal transition”, can be linked to cancer cell motility *p*_migration_. We further link three gene sets to the cell cycle time *t*_proliferation_ and one to the influence of SUP *τ*_SUP_. Another two we link to the CTL recruitment rate *r*_CTL_. and yet another two to the influence of ENH *τ*_ENH_.

### 3.3 Parameter sensitivity analysis indicates multiple mechanisms of therapy resistance

To consider conditions of limited computational power in our analysis that would be found in the analysis of any large-scale ABM, we constrained the global sensitivity analysis to five parameters, excluding the enriched parameter *p*_mutation_, which is the least influential of the selected parameters in the local sensitivity analysis (cf. supplementary Fig. S3). Global sensitivity analysis showed that a large proportion of simulations ended either with an emerging metastasis or complete remission (Fig. 6A). We therefore categorized the labels in emerging metastasis, residual disease and complete remission. The influence of the parameters, either described as partial rank correlation coefficients (prccs) or parameter importance of the trained decision tree is shown in Fig. 6.B/C. Both metrics agree that cancer cell cycle time and motility are more influential than the immune response-related CTL recruitment rate and the cytokine influences. The signs of the prccs follow the intuitive interpretation: long cell cycle times, high CTL recruitment rates and higher influence of ENH tend to lead to better removal of cancer cells, while higher cancer cell motility and higher influence of SUP lead to worse removal.

**Fig. 6.**
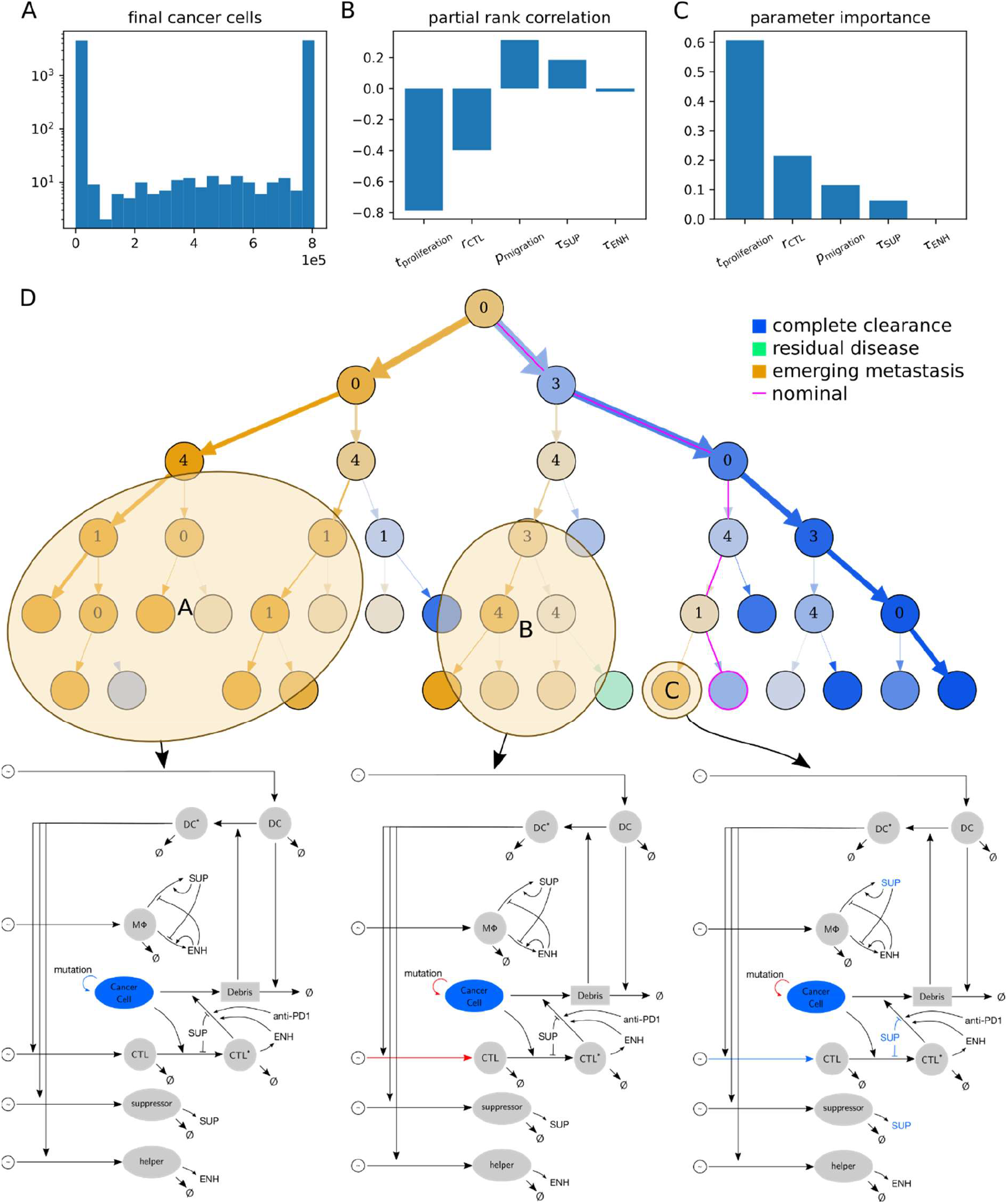
A: Distribution of final cancer cell populations. B: Partial rank correlation coefficients. C: Parameter importances of the decision tree. D: Decision tree for the final cancer cell population. Each node predicts a simulation outcome for a region of the sampled parameter space. At a branch parent node, the parameter space is split along a threshold for a split parameter. The numbers in the parents refer to split parameters: 0: cell cycle time of cancer cells *t*_proliferation_, 1: influence of immunosuppressive cytokines *τ*_SUP_, 3: CTL recruitment rate *r*_CTL_, 4: migration probability of cancer cells *p*_migration_. The color saturation indicates node impurity. Model drafts are shown for highlighted ICI resistant parameter subspaces with color indicating model components influenced by parameter deviations from the nominal configuration (blue: lower/ red: higher values).

The decision tree (Fig. 6.D, supplementary Fig. 4) indicates that, on the one hand the simulated ICI is effective in multiple conditions of the TME, while on the other hand there are different mechanisms of therapy resistance. For the ICI resistant regions, the mechanisms affected by their decision paths are shown colored in the model draft. The tumor tends to be ICI resistant in the following cases: aggressive tumors with short replication time and at least moderate motility, tumors with longer replication time but high motility with smaller CTL recruitment and a case with higher CTL recruitment and longer cell replication time but high influence of SUP and high cancer cell motility. These findings show why it is so difficult to predict ICI response (Morrison et al. 2018), as they indicate that there is a spectrum of different counter-balancing mechanisms that influence its effectiveness.

## 4 Discussion

The aim of this paper is to develop an approach to integrate transcriptomic data into computational models of cell-to-cell interactions in cancer. There is abundant published material about integrating these types of data into unsupervised and supervised machine learning models for the classification and prediction of cancer patient samples. However, to date little has been done regarding merging these data with tissue-level mechanistic computational models, allowing for computer model-supported interpretation of patient data. To this end, we implemented a hybrid, agent-based model describing the interplay between cancer and immune cells in melanoma micrometastasis. To build and characterize the model, we used knowledge of melanoma immunology and publicly available quantitative data describing the behavior of the melanoma cells and different immune cells infiltrating the TME.

There are similar cancer models proposed in the literature. Wang et al. (2013) developed an agent-based melanoma model accounting for cytokine mediated angiogenesis. Hatzikirou et al. (2012) modeled tumor invasion with a lattice gas cellular automaton. Gong et al. (2017) modeled the tumor immune response to PD-1/PD-L1 inhibition. They identified tumor mutational burden and antigen strength as key factors that influence the recruitment of immune cells. They simulate therapy with checkpoint inhibition by changing a model parameter (probability of T cell suppression) at a set time point during a simulation, the same approach we use to model therapy. Compared to the model proposed in this work we do not model CTL proliferation at the TME (cf 2.1.2). Instead, our model considered a greater extend of cell types, including DCs, helper and suppressor cells, and variability in tumor antigens. While this increases the complexity of the model, it hypothetically allows for a more detailed projection of the differential regulation data into the model. The model has some limitations though, which arise from abstractions and simplification, as well as from incomplete knowledge on the cellular mechanisms. For instance, the finite allele mutation model does not replicate the significance of mutational burden on the prognosis of ICI (Morrison et al. 2018).

Here we combined gene set enrichment analysis of cancer immunotherapy response data and global sensitivity analysis of systematic model simulations as a method to constrain the analysis to select model parameters (and connected biological processes) in computational models with large parameter spaces, which cannot be analyzed as a whole due to limitations in available computational power. In our case, a systematic exploration of the entire parameter space would require about 5^28^ simulations, while the application of the method allowed us to reduce the effort to 9,375 simulations and about 175 hours of computation. In contrast to previous approaches to analyze such models, where parameter selection is performed solely hypothesis driven or by requirement as calibration data is missing, our approach offers to perform this in a data driven fashion. Based on our analysis, we hypothesize a causal relationship between given differentially regulated gene sets and cell functions and phenotypes associated with selected cell types in the micrometastasis. In the case of our model, investigation of the model behavior restricted to the selected parameters displayed 3 different mechanistic scenarios of ICI treatment resistance. Key players to these mechanisms are cancer cell motility, which has been previously shown (Dreyer et al. 2018), CTL infiltration, which corresponds to the “warm” vs “cold” tumor hypothesis (Maleki Vareki 2018), and suppressive signaling (TGFβ: Zhao et al. 2018, IDO for non-small-lung-cancer: Botticelli et al. 2018). A limitation of the method is that the linkage between parameters and gene sets remains a manual curation step and depends on the vast expert knowledge of the modelers. Further, it is possible that the enrichment analysis renders gene sets as relevant that are not linked to any parameter in the current instance of the model. In this regard, one can utilize the approach as a method for data-driven, systematic model expansion. For instance, in our analysis three metabolism related genes set are enriched between responders and non-responders, which might encourage to model more details on the cells metabolism and expanding the model with nutrients. This would give the possibility to capture the interplay between immune and metabolic processes.

Furthermore, the linkage as described here is based solely on gene set enrichment analysis and remains a qualitative step, yielding only categorical classification of parameters rather than quantitative differentiation. The latter would enable to deduce parameter perturbations from the data directly, while here we just narrow a selection of parameters, which is subsequently investigated in further detail in a global sensitivity analysis. In this regard, we think that the method can be expanded to generate a quantitative link of transcriptomic data to parameters by calculating a magnitude of the differential regulation and mapping it as a perturbation level to the respective parameters relative to their nominal calibration. This however requires annotation of all parameters with associated gene sets, which poses intensive manual work, but could be supported by automated methods such as text mining.

We think that the combination of enrichment analysis of transcriptomics data and global sensitivity analysis can be applied generally to agent-based or ODE models reflecting cell-to-cell and tissue interactions in cancer and other pathologies (cf. Fig. 3). To this end, it is necessary to have a significant amount of annotated transcriptomics data reflecting the investigated conditions or progression of the disease. When selecting the number of parameters to be explored, one has to consider a trade-off between sufficient sampling of the chosen sub-parameter space and keeping the required computational load in control.

## Supporting information

Supplementary Material for: Integration of transcriptomics data into agent-based models of solid tumor metastasis

## Acknowledgements

The authors thank Martin Eberhardt and Christopher Lischer for useful discussion.

## Data Availability

The code of the model generated for this work is publicly accessible on https://zenodo.org/record/6393283

## Ethics

This article does not contain any studies involving human or animal participants.

## Funding

This work has been supported by the German Ministry of Education and Research (BMBF) through the initiatives e:Med-MelAutim on cancer and autoimmunity [01ZX1905A] and KI-VesD on computation modelling and artificial intelligence guided cancer diagnostics [031L0244A]. We also received funding for our computer model-based research in melanoma from the Manfred Roth-Stiftung, the Trunk-Stiftung and the Matthias-Lackas-Stiftung.

## Conflicts of Interest

The authors declare no potential conflicts of interest.

## Notes

https://zenodo.org/record/6393283

### Competing Interest Statement

The authors have declared no competing interest.

