## Supplementary Material for: Integration of transcriptomics data into agent-based models of solid tumor metastasis for "Integration of transcriptomics data into agent-based models of solid tumor metastasis"

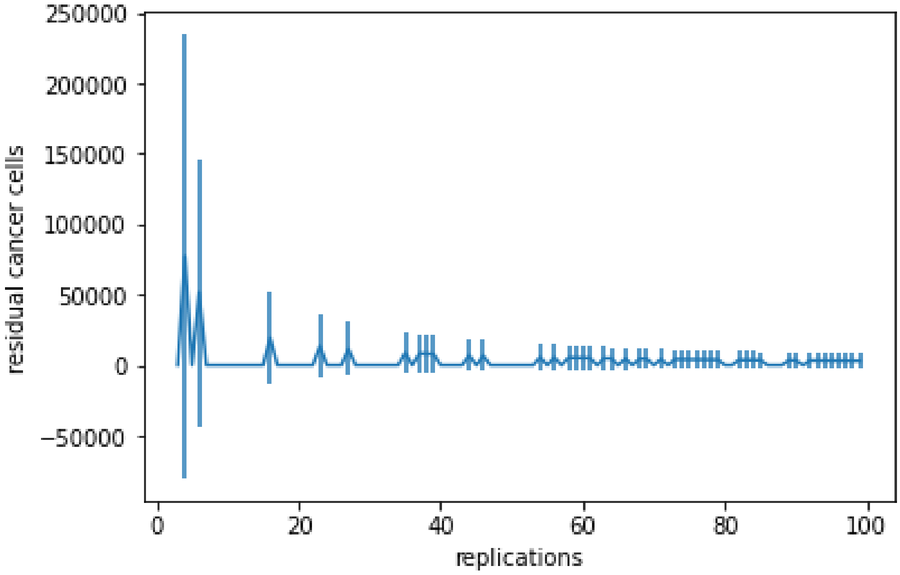


Supplementary figure S1: Estimation of mean cancer cell populations at the end of simulations. The errorbars denote the 95% confidence interval.

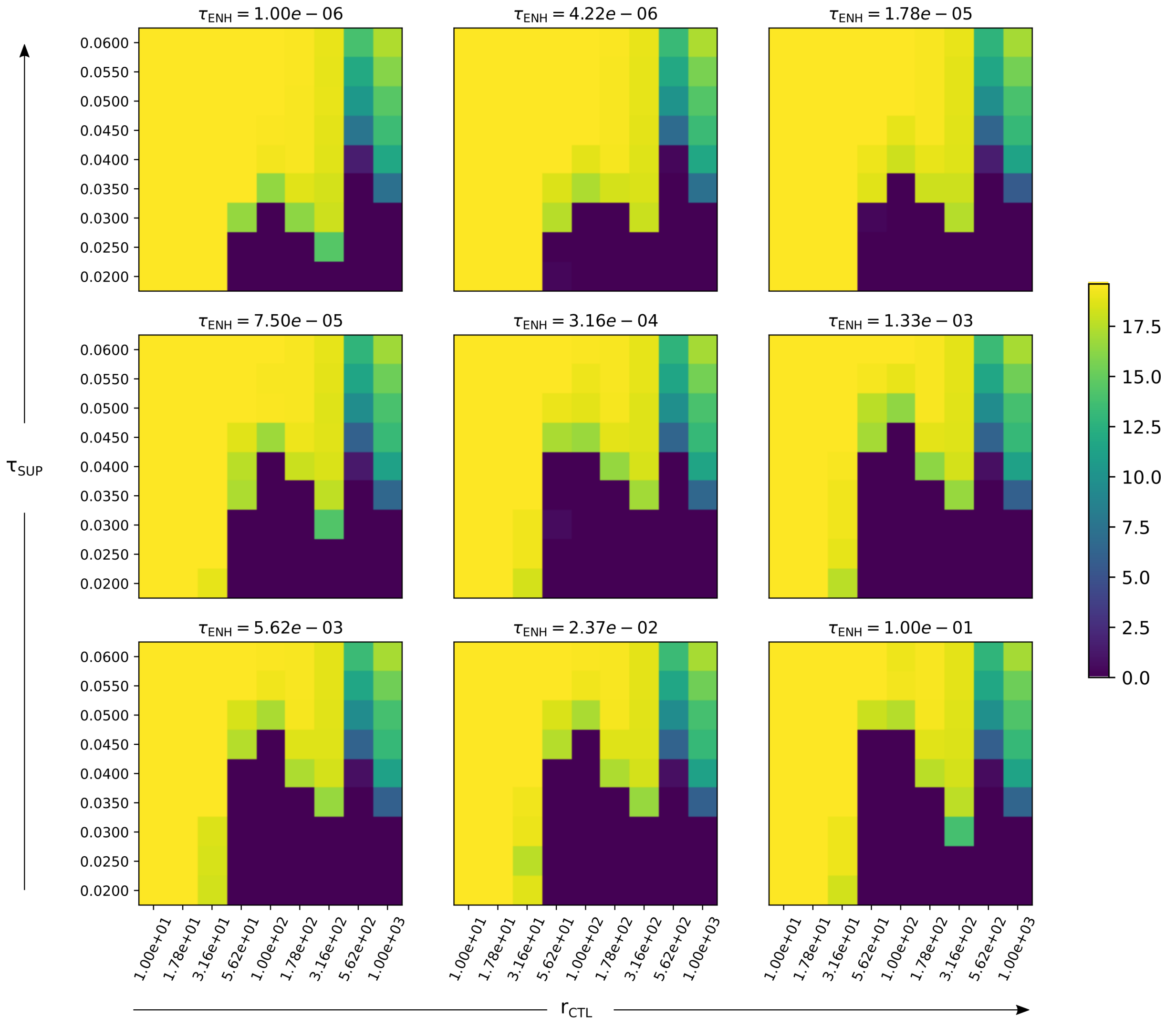


Supplementary figure S2: Parameter space of the uncalibrated parameters. The binary logarithm of the final cancer cell population is color coded.

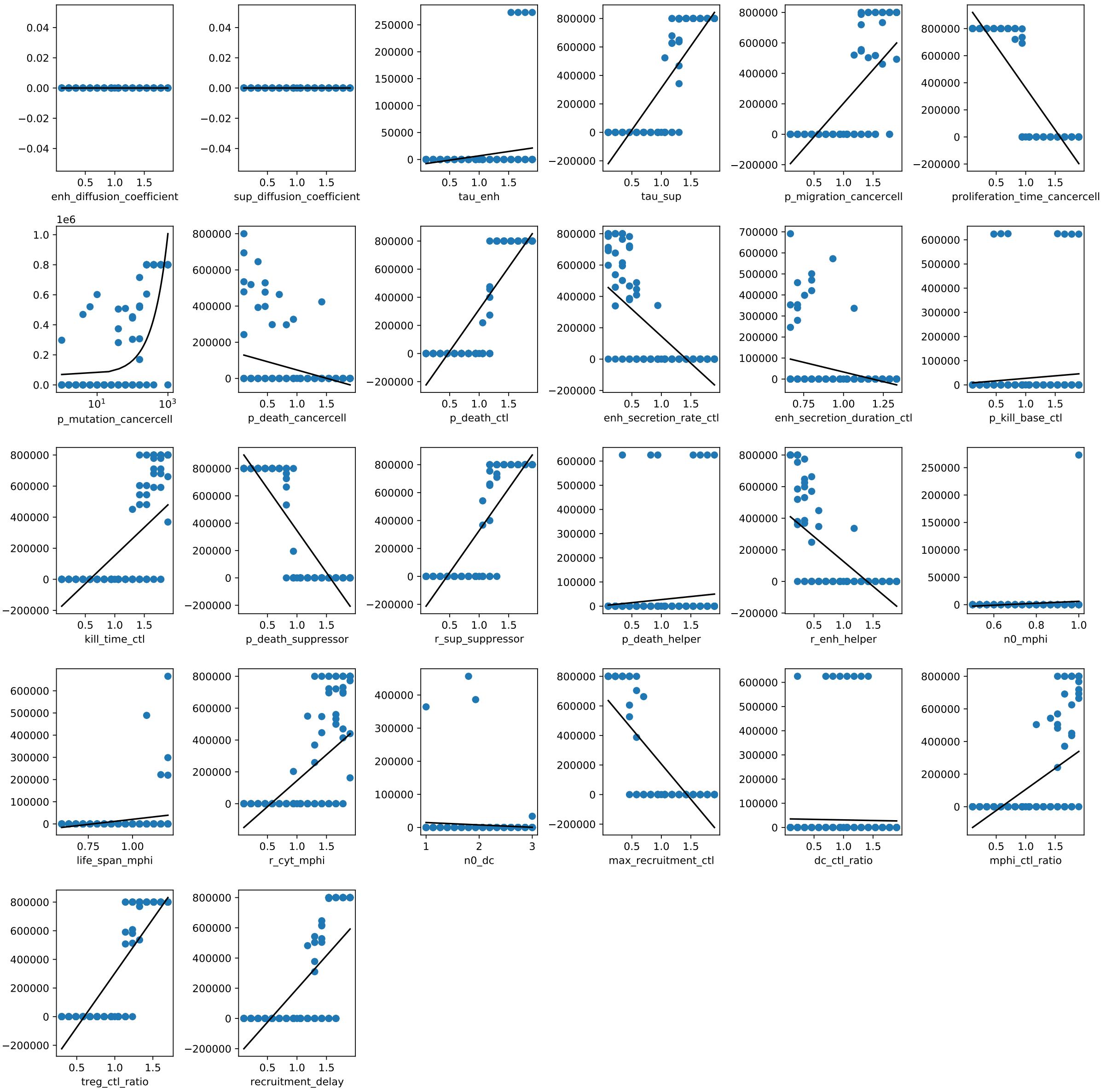


Supplementary figure S3: Local sensitivity analysis of the model parameters. The black lines denote linear regressions.

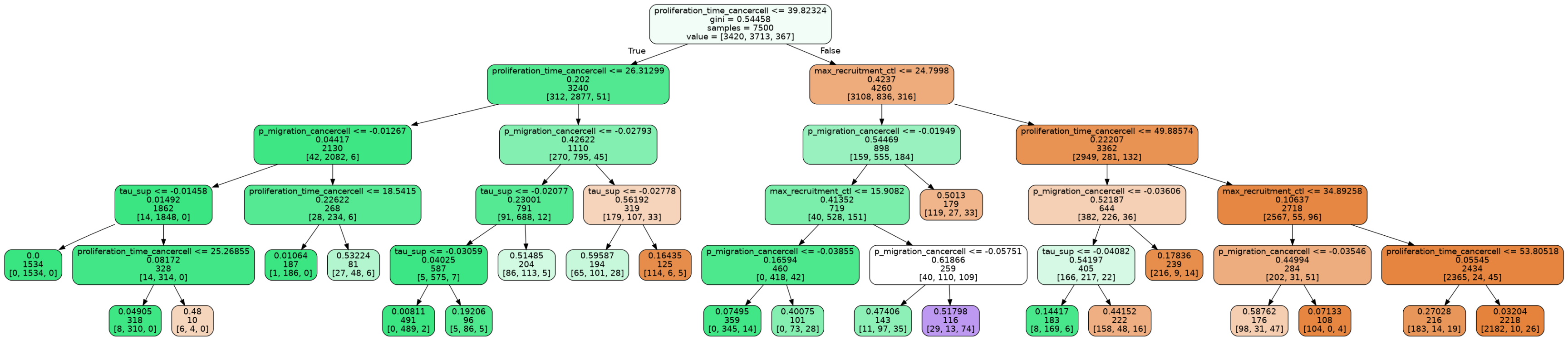


Supplementary figure S4: Decision tree with detailed information on the splits. Values for $\tau_{\text{SUP}}$ and $p_{\text{migration}}$ are negated. The first line of a node contains the split parameter and threshold, the second its gini impurity, the third the number of simulations that are accounted by the node, and the fourth line the number of simulations in the different classes [remission, metastasis, residual disease].

Supplementary table 1: Summary of the model parameters. *Parameter estimation from data in the reference.

| Parameter | Description | Nominal | Interval | Reference |
|---|---|---|---|---|
| Cell framework | | | | |
| $l$ | Cell side length | $10\,\mu m$ | | [9] |
| | Lattice size | $100 \times 100 \times 100 (1\,mm^3)$ | | [9] |
| $\Delta t$ | Time step | $10\,min$ | | [9] |
| Cytokines | | | | |
| $D_{ENH}$ | Diffusion coefficient for ENH | $36\,000\,\mu m^2\,h^{-1}$ | | [3] |
| $D_{SUP}$ | Diffusion coefficient for SUP | $36\,000\,\mu m^2\,h^{-1}$ | | [3] |
| $\tau_{ENH}$ | Influence of ENH on the kill probability | 0.01 | | estimated |
| $\tau_{SUP}$ | Influence of SUP on the kill probability | 0.04 | | estimated |
| Cancer cells | | | | |
| $p_{migration}$ | Moving probability of cancer cells | 0.05 | | [14]* |
| $t_{proliferation}$ | Cell cycle time of cancer cells | $46\,h$ | | [14], [2] |
| $p_{mutation}$ | Mutation probability of cancer cells per gene per cell division | $10^{-6}$ | $[10^{-6}, 10^{-3}]$ | [14], [6] |
| $p_{death}$ | Dying probability of cancer cells | 0.0002 | | [13] |
| CTLs | | | | |
| $p_{death,CTL}$ | Dying probability of CTLs at the TME | 0.0028 | | [5] |
| $r_{ENH}$ | Secretion rate of ENH by CTLs | $6000\,h^{-1}$ | | [11] |
| $t_{ENH}$ | Secretion duration of ENH by CTLs | $3\,h$ | [2, 4] | [4] |
| $p_{kill,base}$ | Base kill probability of CTLs | 1/3 | | [18] |
| $t_{kill}$ | CTL inactivation time after killing a cancer cell | $50\,min$ | | [10] |
| Suppressors | | | | |
| $p_{death,supressor}$ | Dying probability of suppressor cells at the TME | 0.0006 | | [17] |
| $r_{SUP}$ | Secretion rate of SUP by suppressor cells | $6000\,h^{-1}$ | | [11] |
| Helper | | | | |
| $p_{death,helper}$ | Dying probability of helper cells at the TME | 0.0006 | | [17] |
| $r_{ENH}$ | Secretion rate of ENH by helper cells | $6000\,h^{-1}$ | | [11] |
| Macrophages | | | | |
| $n_{0,macrophage}$ | Initial macrophage population size | 500 | [250, 500] | [8] |
| $t_{life,macrophage}$ | Life span of macrophages at the TME | $120\,h$ | [72, 144] | [15] |
| $r_{cytokine}$ | Secretion rate of cytokines by macrophages | $6000\,h^{-1}$ | | [11] |

*Continued on next page*

| Parameter | Description | Nominal | Interval | Reference |
| --- | --- | --- | --- | --- |
| DCs | | | | |
| $n_{0,\mathrm{DC}}$ | Initial DC population size | 1000 | [1000, 3000] | [16] |
| Recruitment | | | | |
| $r_{\mathrm{CTL}}$ | Recruitment rate of CTLs to the TME | $100\,\mathrm{h}^{-1}$ | [10, 1000] | |
| $q_{\mathrm{macrophage}}$ | Macrophage / CTL ratio | $0.37 \cdot r_{\mathrm{CTL}}$ | | [7] |
| $q_{\mathrm{Tcells}}$ | Treg / CTL ratio | 0.076 | $[0.023, 0.129]$ | [12] |
| $t_{\mathrm{delay}}$ | Delay between DC antigen report and CTL recruitment | $168\,\mathrm{h}$ | | [1] |

Supplementary table 2: Enrichment of Hallmark gene sets in anti-PD1 responders vs. non-responders (GSE78220). Significant results (padj $< 0.05$) are annotated with model parameters.

pathway-enrichment-GSE78220

| | pathway | pval | padj | log2err | ES | NES | size | parameter |
|---|---|---|---|---|---|---|---|---|
| 55 | HALLMARK_XENOBIOTIC_METABOLISM | 5.8996765204063E-15 | 1.76990295612189E-13 | 0.996986217533181 | -0.787889543790827 | -2.10692052317677 | 197 | none |
| 5 | Genome.Instability.and.Mutation | 3.93852744784999E-15 | 1.76990295612189E-13 | 0.996986217533181 | -0.779905886562157 | -2.08910623317467 | 216 | p_mutation |
| 15 | HALLMARK_COAGULATION | 4.36961862665697E-14 | 8.73923725331394E-13 | 0.965327754226083 | -0.829290945274524 | -2.15389992515777 | 137 | none |
| 13 | HALLMARK_BILE_ACID_METABOLISM | 2.24270774194028E-09 | 3.36406161291043E-08 | 0.774939030136436 | -0.794792264617605 | -2.0177784415499 | 112 | none |
| 19 | HALLMARK_EPITHELIAL_MESENCHYMAL_TRANSITION | 1.70132189173136E-08 | 2.04158627007763E-07 | 0.73376198835648 | 0.620203208083549 | 1.96579778162797 | 195 | p_migration |
| 57 | Reprogramming.Energy.Metabolism | 4.78266835384818E-08 | 4.78266835384818E-07 | 0.719512826338911 | -0.601719955041389 | -1.68253646374854 | 443 | cell cycle time |
| 4 | Evading.Immune.Destruction | 8.69381860777817E-08 | 7.45184452095272E-07 | 0.704975715167238 | -0.574318497572625 | -1.62650744463243 | 586 | tau_sup |
| 42 | HALLMARK_P53_PATHWAY | 6.29050725153679E-06 | 4.37479724540042E-05 | 0.610526878385931 | -0.653777208541138 | -1.74433538366838 | 192 | none |
| 18 | HALLMARK_E2F_TARGETS | 6.56219586810063E-06 | 4.37479724540042E-05 | 0.610526878385931 | 0.559353076117229 | 1.77292703721374 | 195 | none |
| 22 | HALLMARK_FATTY_ACID_METABOLISM | 1.28601896478468E-05 | 7.71611378870808E-05 | 0.593325476396405 | -0.679397216532026 | -1.78271648070086 | 156 | none |
| 16 | HALLMARK_COMPLEMENT | 2.11001801319693E-05 | 9.73854467629354E-05 | 0.575610261071129 | -0.638595850319474 | -1.70949392149987 | 199 | none |
| 33 | HALLMARK_KRAS_SIGNALING_DN | 2.0700072720978E-05 | 9.73854467629354E-05 | 0.575610261071129 | -0.640927434127702 | -1.70884086590784 | 194 | none |
| 23 | HALLMARK_G2M_CHECKPOINT | 2.05118221669488E-05 | 9.73854467629354E-05 | 0.575610261071129 | 0.531879302736931 | 1.67877896765981 | 190 | cell cycle time |
| 39 | HALLMARK_MYOGENESIS | 8.78796516956346E-05 | 0.000376627078696 | 0.538434096309916 | -0.621660595858179 | -1.66328982753649 | 198 | none |
| 21 | HALLMARK_ESTROGEN_RESPONSE_LATE | 0.000147232276257 | 0.000588929105029 | 0.518848077743792 | -0.614405222180608 | -1.64057837042737 | 196 | none |
| 35 | HALLMARK_MITOTIC_SPINDLE | 0.000182592790313 | 0.000684722963672 | 0.518848077743792 | 0.506421461343078 | 1.60978158781805 | 197 | cell cycle time |
| 52 | HALLMARK_UV_RESPONSE_DN | 0.005291171117345 | 0.01867472159063 | 0.407017918923954 | 0.495074612186953 | 1.50549005953459 | 138 | none |
| 20 | HALLMARK_ESTROGEN_RESPONSE_EARLY | 0.006424522630287 | 0.021415075434291 | 0.407017918923954 | -0.549728582086951 | -1.46912837493279 | 195 | none |
| 60 | Tumor.Promoting.Inflammation | 0.008139296271105 | 0.025703040856121 | 0.380730400722792 | 0.337717233573597 | 1.20100779686102 | 620 | max_recruitment_ctl |
| 10 | HALLMARK_APICAL_JUNCTION | 0.009269061373386 | 0.027807184120157 | 0.380730400722792 | -0.548989604367774 | -1.46387356556655 | 193 | none |
| 50 | HALLMARK_TNFA_SIGNALING_VIA_NFKB | 0.009766593253799 | 0.027904552153711 | 0.380730400722792 | 0.438763070423865 | 1.39554540033859 | 198 | tau_enh |
| 44 | HALLMARK_PEROXISOME | 0.010252803780436 | 0.027962192128463 | 0.380730400722792 | -0.607925333561322 | -1.51927544064341 | 104 | none |
| 30 | HALLMARK_INFLAMMATORY_RESPONSE | 0.010822749461056 | 0.028233259463624 | 0.380730400722792 | 0.4236442901761 | 1.34665457561982 | 197 | max_recruitment_ctl |
| 31 | HALLMARK_INTERFERON_ALPHA_RESPONSE | 0.01640640778502 | 0.041016019462551 | 0.352487857583619 | 0.500859791657303 | 1.44045244582158 | 95 | tau_enh |
| 59 | Sustaining.Proliferative.Signaling | 0.023046092184369 | 0.055310621242485 | 0.30068212900741 | -0.413550259272206 | -1.18823753735596 | 1257 | |
| 36 | HALLMARK_MTORC1_SIGNALING | 0.030283875915986 | 0.069885867498429 | 0.352487857583619 | 0.396431971956246 | 1.25653186065591 | 195 | |
| 32 | HALLMARK_INTERFERON_GAMMA_RESPONSE | 0.03365784430014 | 0.074795209555866 | 0.321775918075361 | 0.411453729173949 | 1.30868433992441 | 198 | |
| 37 | HALLMARK_MYC_TARGETS_V1 | 0.044196982670606 | 0.094707820008441 | 0.321775918075361 | 0.400010705753871 | 1.27090497254369 | 194 | |
| 48 | HALLMARK_SPERMATOGENESIS | 0.074257425742574 | 0.153636053260498 | 0.376039306993639 | 0.406025689572192 | 1.22187751992126 | 132 | |
| 49 | HALLMARK_TGF_BETA_SIGNALING | 0.165562913907285 | 0.32258064516129 | 0.199915231309662 | 0.449105480647152 | 1.18279092541582 | 54 | |
| 54 | HALLMARK_WNT_BETA_CATENIN_SIGNALING | 0.166666666666667 | 0.32258064516129 | 0.186432558434075 | 0.479444305518297 | 1.22632883794447 | 42 | |
| 25 | HALLMARK_HEDGEHOG_SIGNALING | 0.174311926605505 | 0.326797385620915 | 0.127505315300183 | -0.591177997236322 | -1.25965534902625 | 34 | |
| 2 | Enabling.Replicative.Immortality | 0.185185185185185 | 0.326797385620915 | 0.323470511553299 | 0.327673386973156 | 1.09431907785968 | 299 | |
| 29 | HALLMARK_IL6_JAK_STAT3_SIGNALING | 0.184873949579832 | 0.326797385620915 | 0.213927855492356 | 0.414482299518257 | 1.1711138650371 | 87 | |
| 9 | HALLMARK_ANGIOGENESIS | 0.211480362537764 | 0.362537764350453 | 0.11331290842208 | -0.566312286803746 | -1.22069517036301 | 36 | |
| 58 | Resisting.Cell.Death | 0.244979919678715 | 0.408299866131191 | 0.080419996876154 | -0.370715240108163 | -1.06462850675795 | 1145 | |
| 28 | HALLMARK_IL2_STAT5_SIGNALING | 0.30379746835443 | 0.492644543277455 | 0.204294756516886 | 0.331780476331377 | 1.05161230361103 | 195 | |
| 3 | Evading.Growth.Suppressors | 0.339487179487179 | 0.536032388663968 | 0.06494076846129 | -0.370347975765586 | -1.05436860900179 | 661 | |
| 14 | HALLMARK_CHOLESTEROL_HOMEOSTASIS | 0.358208955223881 | 0.551090700344432 | 0.076279718080927 | -0.44899504657103 | -1.07172981411457 | 73 | |
| 11 | HALLMARK_APICAL_SURFACE | 0.453592814371257 | 0.680389221556886 | 0.069283646739255 | -0.457859969692331 | -1.00720146450371 | 43 | |
| 27 | HALLMARK_HYPOXIA | 0.488649940262843 | 0.698071343232634 | 0.054907370329096 | -0.370898757573757 | -0.988211935132508 | 191 | |
| 40 | HALLMARK_NOTCH_SIGNALING | 0.485875706214689 | 0.698071343232634 | 0.10027910675413 | 0.415527621832798 | 0.995274280728132 | 32 | |
| 34 | HALLMARK_KRAS_SIGNALING_UP | 0.541120381406436 | 0.755051694985725 | 0.050092287440211 | -0.36101694421271 | -0.962646937482929 | 193 | |
| 56 | Inducing.Angiogenesis | 0.576759061833689 | 0.776447105788423 | 0.042077210988939 | -0.344817717588742 | -0.968023769925494 | 485 | |
| 43 | HALLMARK_PANCREAS_BETA_CELLS | 0.582335329341317 | 0.776447105788423 | 0.057246113520288 | -0.425975218704307 | -0.926972169304473 | 39 | |
| 26 | HALLMARK_HEME_METABOLISM | 0.605952380952381 | 0.790372670807453 | 0.044864511316856 | -0.347290558938431 | -0.926601910338933 | 192 | |
| 7 | HALLMARK_ALLOGRAFT_REJECTION | 0.720379146919431 | 0.919632953514168 | 0.036685042761344 | -0.326948822430929 | -0.873758083235641 | 195 | |
| 45 | HALLMARK_PI3K_AKT_MTOR_SIGNALING | 0.784105960264901 | 0.972145615002758 | 0.03792714671081 | -0.330190176990654 | -0.825183289705759 | 104 | |
| 38 | HALLMARK_MYC_TARGETS_V2 | 0.793918918918919 | 0.972145615002758 | 0.08243440915271 | 0.306867361044419 | 0.828571720318353 | 58 | |
| 12 | HALLMARK_APOPTOSIS | 0.821621621621622 | 0.985945945945946 | 0.107972360317345 | 0.289628063139072 | 0.894820462675071 | 159 | |
| 1 | Activating.Invasion.and.Metastasis | 0.887210473313192 | 1 | 0.016838053046391 | -0.305028153685303 | -0.875143862528585 | 1102 | |
| 6 | HALLMARK_ADIPOGENESIS | 0.896428571428571 | 1 | 0.02618415159462 | -0.294301874693068 | -0.785223416785341 | 192 | |
| 53 | HALLMARK_UV_RESPONSE_UP | 0.906288532675709 | 1 | 0.027448008180102 | -0.29234860415355 | -0.767653307450578 | 154 | |
| 24 | HALLMARK_GLYCOLYSIS | 0.938315539739027 | 1 | 0.023508484501329 | -0.283691784971784 | -0.758629237720138 | 197 | |
| 41 | HALLMARK_OXIDATIVE_PHOSPHORYLATION | 0.984468339307049 | 1 | 0.021145636251359 | -0.267145829551459 | -0.711018394854852 | 184 | |
| 46 | HALLMARK_PROTEIN_SECRETION | 0.969855832241153 | 1 | 0.02709674231833 | -0.277099010616261 | -0.686106707802994 | 94 | |
| 8 | HALLMARK_ANDROGEN_RESPONSE | 0.993394980184941 | 1 | 0.026254339649827 | -0.255947489501375 | -0.633525127498473 | 96 | |
| 17 | HALLMARK_DNA_REPAIR | 1 | 1 | 0.022726926034007 | -0.153287435758237 | -0.400531437285294 | 147 | |
| 47 | HALLMARK_REACTIVE_OXYGEN_SPECIES_PATHWAY | 1 | 1 | 0.066896627650067 | 0.167076110181607 | 0.431288020210968 | 47 | |
| 51 | HALLMARK_UNFOLDED_PROTEIN_RESPONSE | 0.905172413793103 | 1 | 0.08862611332851 | 0.276124013534668 | 0.811302214045898 | 107 | |
